# Type I Interferon Signaling on Antigen-Presenting Cells Blunts Cell-Mediated Immunity Toward *Listeria monocytogenes*

**DOI:** 10.1101/2022.12.02.518944

**Authors:** Zachary T. Morrow, John-Demian Sauer

**Affiliations:** Department of Medical Microbiology and Immunology, University of Wisconsin-Madison, Madison WI 53702

**Keywords:** *Listeria monocytogenes*, interferon, inflammasome, vaccine, antigen-presenting-cell

## Abstract

*Listeria monocytogenes* is a facultative intracellular pathogen that has been used for decades to understand mechanisms of bacterial pathogenesis and both innate and adaptive immunity. *L. monocytogenes* is a potent activator of CD8^+^ T-cell mediated immunity. Yet how the innate immune response to infection modulates CD8^+^ T-cell responses is incompletely understood. Here, we utilize an attenuated *L. monocytogenes* vaccine platform to understand the impact of two innate immune pathways, type I interferon and inflammasomes, on CD8^+^ T-cell responses using a combination of mutant mice and genetically engineered *L. monocytogenes*. IFNAR^−/−^ mice had the most robust T-cell response, while Caspase-1^−/−^ mice were not different from WT. We uncover a role for inflammasomes in the absence of type I interferon as Caspase-1^−/−^/IFNAR^−/−^ mice had fewer T-cells than IFNAR^−/−^. IFNAR^−/−^ had more than twice as many memory precursors, promoting enhanced protection from rechallenge. Importantly, increased memory precursor T-cell abundance did not come at the expense of short-lived effectors. Vaccines genetically modified to induce lower type I interferon production yielded enhanced T-cell responses. Deficits from type I interferon signaling are dendritic cell-intrinsic, rather than acting on T-cells, as IFNAR^−/−^ dendritic cells induced two-fold more T-cell proliferation than WT in *ex vivo* T-cell proliferation assays. Thus, modulating type I IFN signaling during vaccination may lead to more potent T-cell-based vaccines. Importantly, this suggests innate immune signaling significantly impacts the CD8^+^ T-cell response and suggests CD8^+^ T-cell quantity and quality are important factors to consider during rational vaccine design.

## Introduction

*Listeria monocytogenes* is a genetically tractable, gram-positive, facultative intracellular bacterium that has been used extensively to study various aspects of cell biology, bacterial pathogenesis, and both innate and adaptive immune responses^1,2^. Due to its ability to stimulate a robust CD8^+^ T-cell response, *L. monocytogenes* has been instrumental in understanding CD8^+^ T-cell responses and has been explored as an anti-cancer vaccine platform^3,4^. Despite years of research, many questions remain about how *L. monocytogenes* triggers robust CD8^+^ T-cell responses and how to rationally design vaccines to generate large quantities of highly functional, long-lived CD8^+^ T-cells.

*L. monocytogenes* that fail to access the host cell cytosol do not induce robust CD8^+^ T-cell responses^5^. This led to the hypothesis that activation of *cytosol-specific* innate immune signaling pathways is critical for the CD8^+^ T-cell response. *L. monocytogenes* activates numerous cytosol-specific innate immune signaling pathways including Nucleotide Oligomerization Domain (NOD)-like receptors (NLRs), the production of lipid mediators of inflammation known as eicosanoids, STING-dependent type I interferon (IFN) production (comprised of IFN-β and the IFN-α family), and inflammasomes^4,6^. Contrary to this hypothesis, many of these pathways proved to be not only dispensable such as TLRs and NOD1/2^6–9^, but in some cases actively detrimental to the T-cell response toward *L.* monocytogenes^6–9^.

Counterintuitively, cytosolic induction of type I IFN negatively impacts *L. monocytogenes* stimulated T-cell responses^4,10–14^. Type I IFN production is triggered by *L. monocytogenes*’ secretion of cyclic-di-AMP (CDA) through multidrug resistance pumps (MDRs) into the host cell cytosol and through cyclic GMP–AMP (cGAMP) synthase (cGAS)-mediated production of cGAMP following recognition of bacterial DNA^15,16^. CDA or cGAMP (collectively referred to as cyclic-di-nucleotides [CDNs]) are recognized by ER bound Stimulator of Interferon Genes (STING)^16^, leading to type I IFN production in a pathway involving TANK Binding Kinase 1(TBK1) and Interferon Regulatory Factor (IRF)3 and IRF7^17^. Mice lacking IRF3/7 (which are necessary for type I IFN production) have an enhanced T-cell response toward *L. monocytogenes*. At the same time, exogenous supplementation of CDNs blunts T-cell responses^14^, collectively indicating that type I IFN is detrimental to T-cell immunity toward *L. monocytogenes*.

Inflammasomes are multiprotein signaling complexes composed of a receptor, the adaptor, Apoptosis-associated Spec-like protein containing a Caspase recruitment domain (ASC), and Caspase-1^12^. Although many inflammasome receptors exist, ligand binding results in activation of the inflammasome effector, Caspase-1, leading to secretion of IL-1β and IL-18 and a Gasdermin-D-dependent inflammatory form of cell death called pyroptosis^18^. *L. monocytogenes* naturally triggers low levels of inflammasome activation primarily through infrequent bacteriolysis in the cytosol that leads to activation of the DNA sensing AIM2 inflammasome^19^. We recently demonstrated that hyperactivation of the inflammasome is detrimental to *Listeria-*stimulated CD8^+^ T-cell responses^13^. How type I IFN or inflammasome activation impairs *L. monocytogenes*-induced T-cell immunity is unknown. Additionally, whether dual blockade of type I IFN and inflammasome activation could additively or even synergistically improve CD8^+^ T-cell responses to *L. monocytogenes* vaccination is unknown.

To test the hypothesis that blocking both type I IFN signaling and inflammasomes would synergistically enhance the T-cell response toward *L. monocytogenes,* we crossed mice lacking the type I IFN receptor (IFNAR^−/−^) and the inflammasome effector Caspase-1 (Caspase-1^−/−^) to generate mice lacking both (Caspase-1^−/−^/IFNAR^−/−^). Then we assessed the primary and recall T-cell responses and protective immunity. Caspase-1^−/−^ mice had no deficits in the T-cell response, likely due to the relatively low-level inflammasome activation induced by *L. monocytogenes*^19^. On the other hand, IFNAR^−/−^ mice developed more robust primary and memory T-cell responses. We uncover a role for inflammasome activation only in the absence of type I IFN signaling as Caspase-1^−/−^/IFNAR^−/−^ mice have reduced T-cell responses compared to IFNAR^−/−^ mice. Using both IFNAR^−/−^ mice as well as *L. monocytogenes* engineered to alter STING-dependent type I IFN production, we demonstrate that type I IFN inhibits memory precursor T-cell formation. Finally, *ex vivo* T-cell proliferation assays demonstrate that type I IFN signaling negatively impacts dendritic cells rather than acting directly on T-cells.

Taken together, this data suggests that inhibiting type I IFN signaling during vaccine application is a rational approach to not only drive numerically more T-cells but promote a more durable response through increased memory-precursor formation and subsequent memory T-cell persistence. Furthermore, identifying that type I IFN specifically impairs APCs may drive more rational vaccine and immunotherapy designs through blocking the detrimental impacts of type I IFN on APCs while sparing the beneficial effects reported on T-cells. Collectively, this suggests a more thorough understanding of innate immune signaling on cell-specific subsets may further enhance rational vaccine design.

## Results

Type I Interferon Impairs the Primary T-cell Response toward *L. monocytogenes* while Inflammasome Activation Plays a Minimal Role

Both type I IFN and inflammasome activation negatively impact the CD8^+^ T-cell response toward *L. monocytogenes*. Mice lacking genes necessary for type I IFN production during *L. monocytogenes* infection (i.e., STING and IRF3/7) develop enhanced cell-mediated immunity and greater recall CD8^+^ T-cell responses^14^ and inflammasome hyperactivation impairs primary CD8^+^ T-cell responses and protective immunity^13^. We, therefore, hypothesized that mice lacking both Caspase-1 and IFNAR would develop synergistically enhanced CD8^+^ T-cell responses compared to the loss of either innate immune pathway alone.

To test this hypothesis, we immunized WT, Caspase-1^−/−^, IFNAR^−/−^ or Caspase-1^−/−^/IFNAR^−/−^ mice with 10^3^ colony-forming units (CFUs) of our attenuated vaccine strain of *L. monocytogenes* (termed LADD) expressing the model antigens B8R and OVA^20,21^. At the peak of the T-cell response, seven days post-immunization, splenocytes were analyzed for IFNγ, TNFα, and IL-2 expression by flow cytometry after *ex vivo* peptide stimulation (S1D). In contrast to our hypothesis that Caspase-1^−/−^ would develop enhanced T-cell responses, Caspase-1^−/−^ and WT mice had similar antigen-specific T-cell frequencies (Fig. 1A, S1A). In agreement with previous studies using STING^−/−^ or IRF3/7^−/−^ deficient mice^14^ suggesting type I IFN signaling impairs T-cell responses but could not rule out STING- or IRF3/7-dependent but type I IFN-independent effects, IFNAR^−/−^ mice developed two-fold more antigen-specific CD8^+^ T-cells (Fig. 1A, S1A). IFNγ expression (measured by MFI) was increased in antigen-specific INFAR^−/−^ CD8^+^ T-cells compared to WT or Caspase-1^−/−^ possibly indicating enhanced function (Fig. 1C, S1C). Despite lacking type I IFN signaling, Caspase-1^−/−^/IFNAR^−/−^ mice developed fewer IFNγ^+^ CD8^+^ cells compared to IFNAR^−/−^ mice (Fig. 1A, S1A), suggesting a beneficial role for inflammasome activation in the absence of type I IFN. However, IFNγ expression trended higher in these mice compared to WT or Caspase-1^−/−^ (Fig. 1C, S1C). IFNAR^−/−^ mice had a greater multifunctional CD8^+^ T-cell response (defined as IFNγ^+^TNFα^+^IL-2^+^) compared to WT, Caspase-1^−/−^, or Caspase-1^−/−^/IFNAR^−/−^ (Fig. 1B, S1B). Taken together our data suggest that type I IFN signaling is detrimental to the primary T-cell response and although inflammasome signaling does not independently contribute to the T-cell response, it is necessary for optimal T-cell responses in the absence of type I IFN signaling.

**Figure 1 –.**
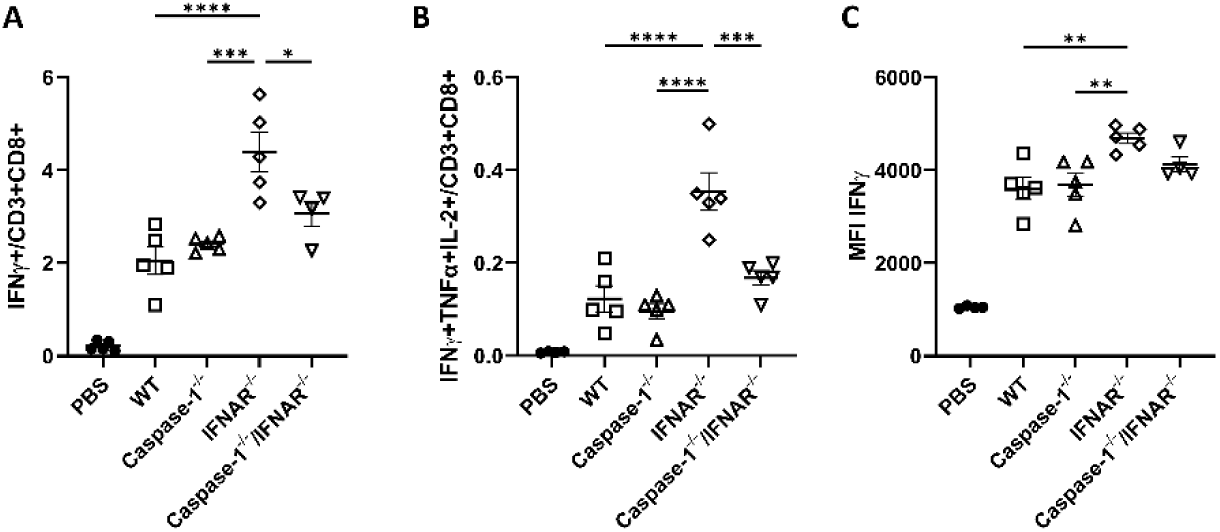
Inflammasome activation and Type I IFN production alter the CD8+ T-cell response to *L. monocytogenes* vaccination. Mice were immunized with 10^3^ LADD expressing B8R and OVA. Splenocytes were examined for OVA specific antigen responses by *ex vivo* peptide stimulation 7 days after immunization. Cells were gated for CD3+ and CD8+ followed by A) IFNγ or B) IFNγ+ TNFα+ and IL-2+ as well as C) IFNγ expression by MFI. Data are representative of two independent experiments of 4-5 mice per group Significance was determined by a one-way ANOVA with Tukey’s correction. * p < 0.05.

We were puzzled that Caspase-1^−/−^ mice did not have improved T-cell responses and hypothesized that impaired inflammasome activation may promote type I IFN production leading to an impairment of an otherwise augmented T-cell response. Indeed, Caspase-1^−/−^ mice produce increased type I IFN production during infection with another intracellular bacterium, *Francisella tulerensis*^22^. To test this hypothesis, we generated bone-marrow-derived macrophages (BMDMs) from each mouse line and infected them with our vaccine strain and assayed the supernatants for either IFN-β or IL-1β as a readout of inflammasome activation. Caspase-1 deficiency potentiated type I IFN production *ex vivo* and IFNAR^−/−^ BMDMs also produced slightly more IFN-β than WT (S2A). Caspase-1 deficiency resulted in complete loss of IL-1β production (Figure S2B) and IFNAR^−/−^ BMDMs produced less IL-1β compared to WT (S1B) suggesting type I IFN may promote inflammasome activation. Together, this data supports a model where blocking inflammasome activation would enhance T-cell responses but increased type I IFN abrogates such enhancement. It remains unclear what inflammasome-dependent signaling is necessary in the absence of type I IFN signaling.

### Type I IFN inhibits memory precursor CD8^+^ T-cell formation in the primary immune response

Our data indicated that blocking type I IFN signaling increased the frequency of multifunctional IFNγ, TNFα and IL-2 producing antigen-specific T-cells. Since multifunctional T-cells may give rise to memory T-cell populations^23–26^, we hypothesized that type I IFN inhibits the formation of memory precursor (MPEC) cells that ultimately give rise to long-term memory CD8^+^ T-cells. MPECs have CD127^high^KLRG1^low/high^ expression, while short-lived effectors (SLECs) have CD127^low^KLRG1^high^ expression^26–29^ (Fig. S4). To test this hypothesis, we examined both the absolute number and relative frequency of OVA- and B8R-tetramer+ MPECs and SLECs in each mouse genotype. Despite differences in tetramer-positive CD8^+^ T-cells (Fig. 2A, D) SLEC numbers were equivalent independent of innate immune signaling (Figure 2B, E). IFNAR^−/−^ mice developed nearly 10-fold more MPECs than WT and Caspase-1^−/−^/IFNAR^−/−^ mice trended toward having more MPECs than WT (Fig. 2C, F). Although there are differences in the frequency of tetramer-positive CD8^+^ T-cells in Caspase-1^−/−^/IFNAR^−/−^mice compared to WT, there were no statistically significant differences in the absolute number per mouse spleen suggesting that inflammasome activation may have little or no role in T-cell fate determination during *L. monocytogenes* immunization at the primary stage. Taken together, our data suggest that MPEC formation is impaired by type I IFN during *L. monocytogenes* vaccination.

**Figure 2 –.**
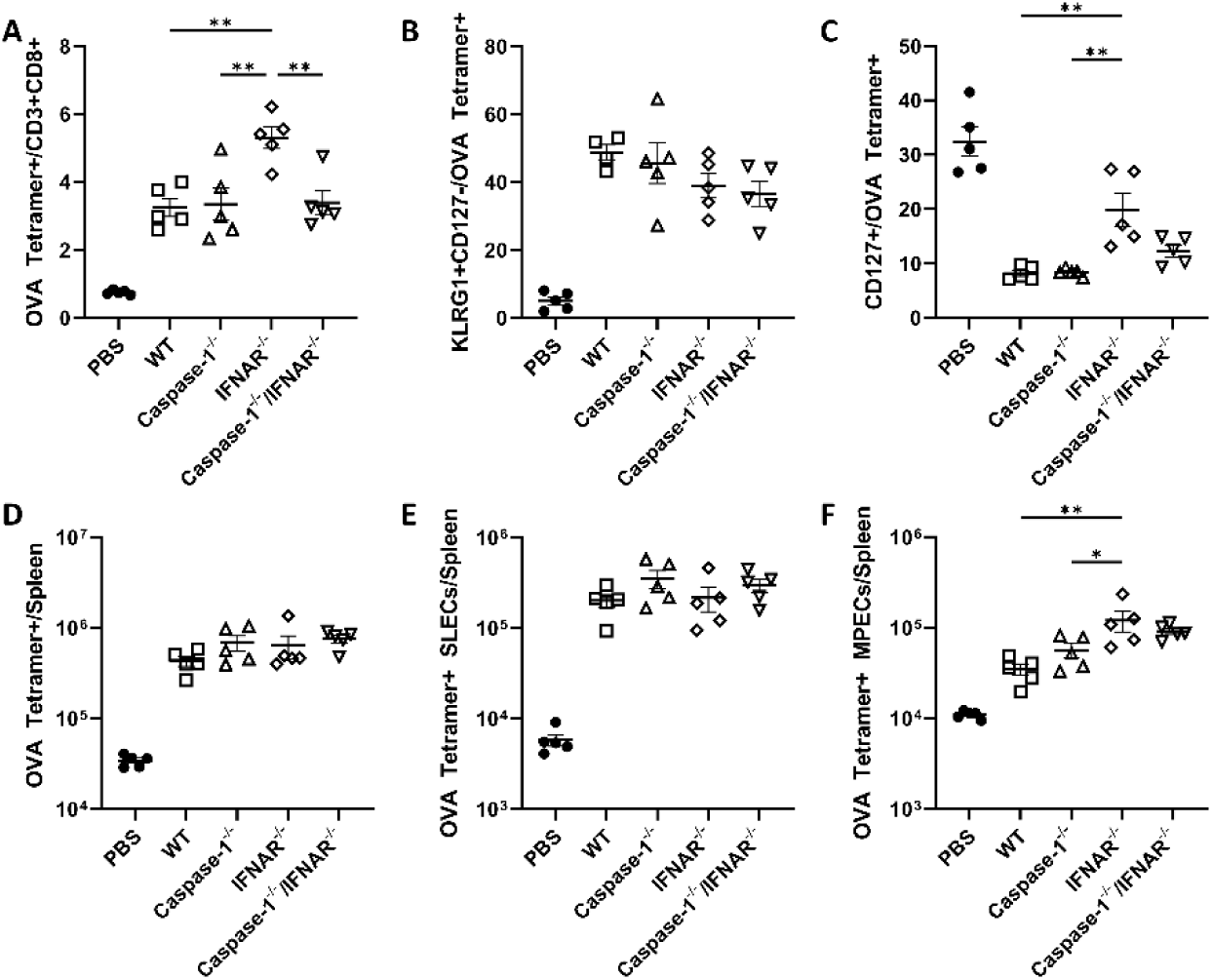
The primary T-cell response is augmented and skewed toward a memory phenotype In the absence of type I IFN. Mice were immunized with10^3^ cfu *L. monocytogenes* expressing B8R and OVA. OVA-specific responses were examined at day 7 post immunization by gating for CD3+ CD8+ and B220-cells and examing A, D) OVA-tetramer+ or B, E) OVA-tetramer+ and KLRG1+ (SLECs) or C, F) OVA-tetramer+CD127+ cells (MPECs). Data shown are representative of two independent experiments of 4-5 mice per group. Significance was determined by a one-way ANOVA with Tukey’s correction. * p < 0.05.

### Augmented T-cell Frequencies Promote Enhanced Vaccine Efficacy and Confer Protection from Rechallenge

We hypothesized that CD8^+^ T-cells would be maintained at greater frequencies in mice lacking type I IFN and this would confer greater protection from rechallenge. To test this hypothesis, we immunized mice with 10^3^ LADD expressing B8R and OVA and waited thirty days after contraction of the primary T-cell response has occured^30^ and examined the memory pool of antigen-specific T-cells that remained. Consistent with the primary T-cell responses and MPEC numbers, WT and Caspase-1^−/−^ mice had similar memory T-cell frequencies while IFNAR^−/−^ mice contained more than double the frequency of CD8^+^ T-cells (Figure 3A, S5A). Moreover, IFNγ expression was higher in IFNAR^−/−^ mice (Fig. S3C, D). Despite trending toward increased frequencies of MPECs and multifunctional T-cells during the primary response, Caspase-1^−/−^/IFNAR^−/−^ mice had comparable CD8^+^ T-cell numbers to WT in this phase of the T-cell response (Fig. 3A, B, S3A, B). These trends were true for multifunctional cells as well (Figure 3B, S3B) and suggest that while loss of type I IFN promotes memory T-cell formation, inflammasome activation may be required for their long-term maintenance in the absence of type I IFN signaling.

**Figure 3 -.**
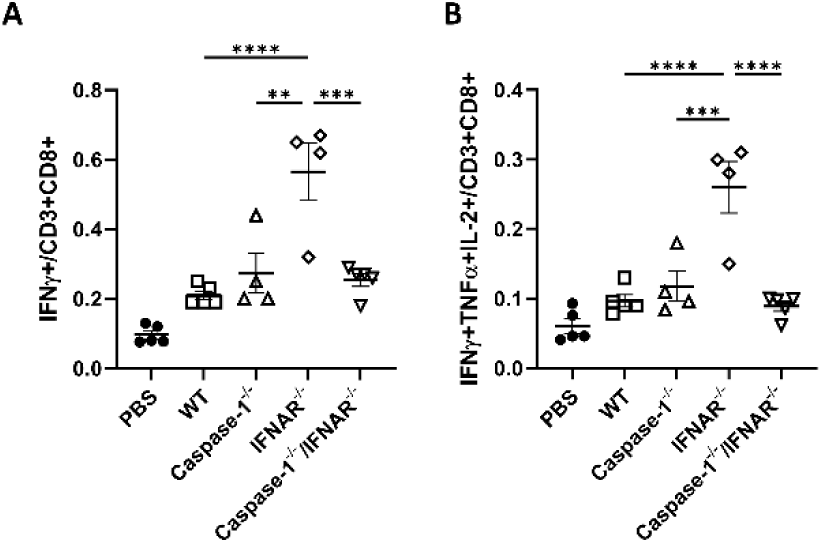
Memory Cells Persist in the absence of type I IFN. Mice were immunized with 1×10^3^ cfu *L. monocytogenes* expressing B8R and OVA. At day 30 post immunization, after contraction is complete, OVA-specific responses were examined by *ex vivo* peptide stimulation followed by intracellular cytokine staining for A) IFNγ producing CD8+ T-cells, or B) IFNγ+TNFα+IL-2+ multifunctional CD8+ T-cells. Data shown are representative of two independent experiments of 4-5 mice per group. Significance was determined by a one-way ANOVA with Tukey’s correction. * p < 0.05.

Increased MPEC numbers in the primary response and memory T-cells thirty days post-immunization in IFNAR^−/−^ indicated that recall responses may be augmented. To test this hypothesis, we immunized mice with 10^3^ LADD expressing B8R and OVA and thirty days later, rechallenged them with 5 LD50s (5 x 10^5^) of WT *L. monocytogenes* expressing B8R and OVA—a dose that would be rapidly lethal to unimmunized mice. Three days-post challenge, we assessed splenocytes for antigen-specific responses as before. Both INFAR^−/−^ and Caspase-1^−/−^/IFNAR^−/−^ mice had twice as many antigen-specific CD8^+^ T-cells as WT and Caspase-1^−/−^ mice upon rechallenge (Fig. 4A, S3A), IFNAR^−/−^ mice had a slightly higher, albeit not statistically significant, frequency of antigen-specific T-cells than Caspase-1^−/−^/IFNAR^−/−^ mice (Figure 4A, S3A). IFNAR^−/−^ CD8^+^ antigen-specific T-cells also expressed higher IFNγ possibly indicating greater functionality (Fig. S6C, D). Taken together this data suggests that the larger pool of MPECs and subsequently larger pool of memory CD8^+^ T-cells in IFNAR^−/−^ mice promote enhanced T-cell recall responses.

**Figure 4 -.**
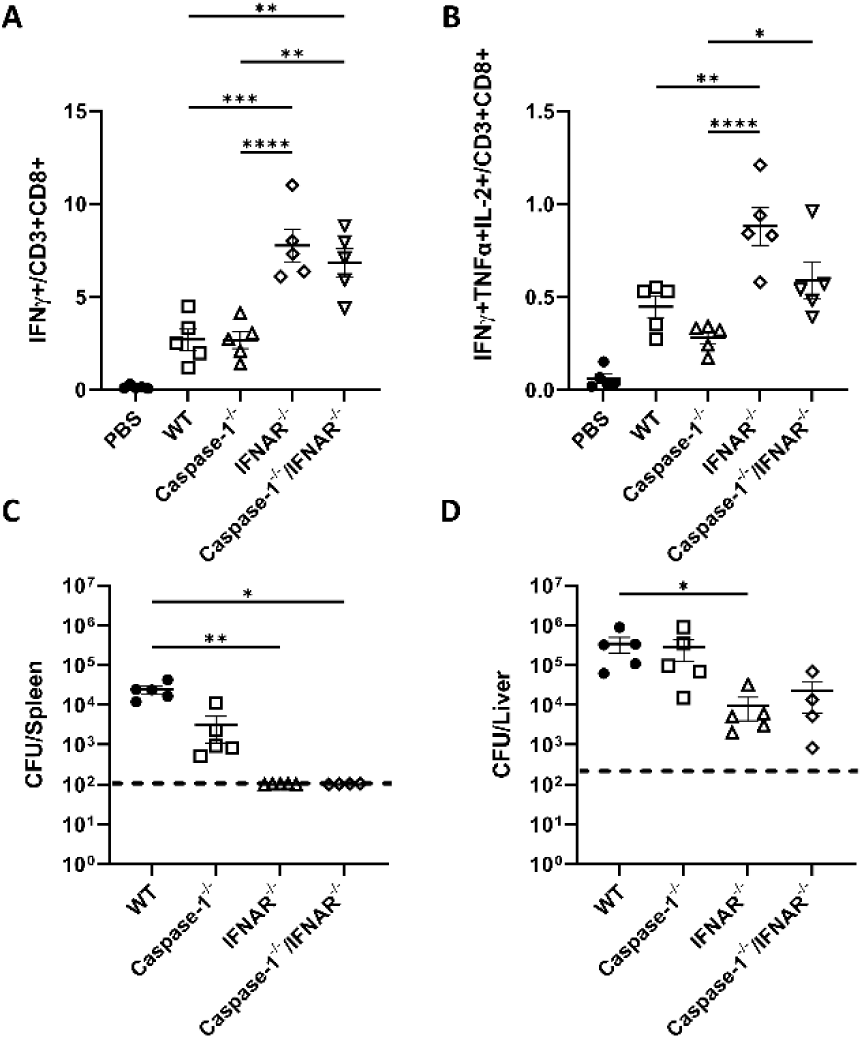
The Recall Response is Enhanced in the absence of type I IFN mice and Promotes Augmented T-cell Memory and Protection. Mice were immunized with 1×10^3^ *L. monocytogenes* expressing B8R and OVA and then challenged with 5×10^5^ WT *L. monocytogenes* expressing B8R and OVA. OVA-specific responses were examined by peptide stimulation at day 3 post challenge for CD8+ T-cells producing A) IFNγ, and B) IFNγ+TNFα+IL-2+ multifunctional T-cells. Additionally, spleens and livers were collected and homogenized and plated for CFU in C) Spleen E) Liver as a measure of protection. Data shown are representative of two independent experiments of 4-5 mice per group. Significance was determined by a one-way ANOVA with Bonferroni’s correction. * p < 0.05, **p <0.01.

Since mice lacking type I IFN signaling have augmented recall responses, we hypothesized that these mice would be better protected from rechallenge indicating increased vaccine efficacy. To test this hypothesis, we immunized and rechallenged mice as before and enumerated bacterial burdens in the liver and spleen as a proxy for vaccine efficacy. Following rechallenge, IFNAR^−/−^ and Caspase-1^−/−^/IFNAR^−/−^ mice had no detectable CFU in the spleen (Figure 4C), whereas WT and Caspase-1^−/−^ mice had detectable bacterial burdens with Caspase-1^−/−^ trending toward less than WT (Fig 4C). In the liver, only IFNAR^−/−^ was significantly protected compared to WT, however, Caspase-1^−/−^/IFNAR^−/−^ mice trended toward lower burdens than WT. Taken together, our data thus far indicate that early priming events may dictate the overall recall response and thus protective immunity. In other words, type I IFN impairs the primary T-cell response, blunting memory T-cell formation, thus impairing recall responses and protective immunity.

### *L. monocytogenes* Engineered to hypo-induce type I IFN Augment the T-cell Response but not Protective Immunity

Since type I IFN is detrimental to T-cell responses and protective immunity, we hypothesized that *L. monocytogenes* engineered to induce less type I IFN would promote enhanced T-cell responses in WT mice and thus represent optimized vaccine platforms. To test this hypothesis, we constructed vaccine strains of *L. monocytogenes* expressing B8R and OVA that hyper- or hypo-induce type I IFN production and assessed the T-cell response. We utilized the previously described *Δmdr-MTAC* and *ΔtetR* strains, which induce less type I IFN or more type I IFN through more or less cyclic-di-AMP secretion, respectively^31,32^. Both strains were assessed for IFN-β production in BMDMs. *Δmdr-MTAC* induced less IFN-β compared to WT, but still more than in uninfected controls which were at the detection limit (Fig. S7). Consistent with previous data^32^, *ΔtetR* mutants induced more type I IFN WT (Fig. S7).

We immunized mice with a low (10^3^) or high (10^7^) dose of the indicated strain and assessed T-cell responses by flow cytometry as before. *ΔtetR* resulted in a 50% reduction in antigen-specific T-cells compared to WT at either dose (Figure 5A, D, S8A, and S9A, D). *Δmdr-MTAC* induced two-fold more antigen-specific T-cells than WT at the low dose (Figure 5A, D, S8A, and S9A, D). However, at the high dose, there was no difference between LADD and *Δmdr-MTAC* suggesting that the residual type I IFN induced by this strain may be sufficient to reduce the T-cell response to WT levels (Fig. 5A, D, S8A, and S9A, D). *Δmdr-MTAC* induced both more SLECs and MPECs than WT or *ΔtetR* (Fig 5B-C, E-F, and S9B-C, E-F). IFNγ expression was largely similar across all strains indicating that even modest levels of type I IFN are sufficient to abrogate the increased IFNγ expression observed in IFNAR^−/−^ mice (Figure S8 C, F). Taken together, our data demonstrate that *L. monocytogenes* can be modified to decrease type I IFN and improve T-cell priming while increasing type I IFN production inhibits *L. monocytogenes*-primed T-cell responses, consistent with previous results^14^.

**Figure 5 -.**
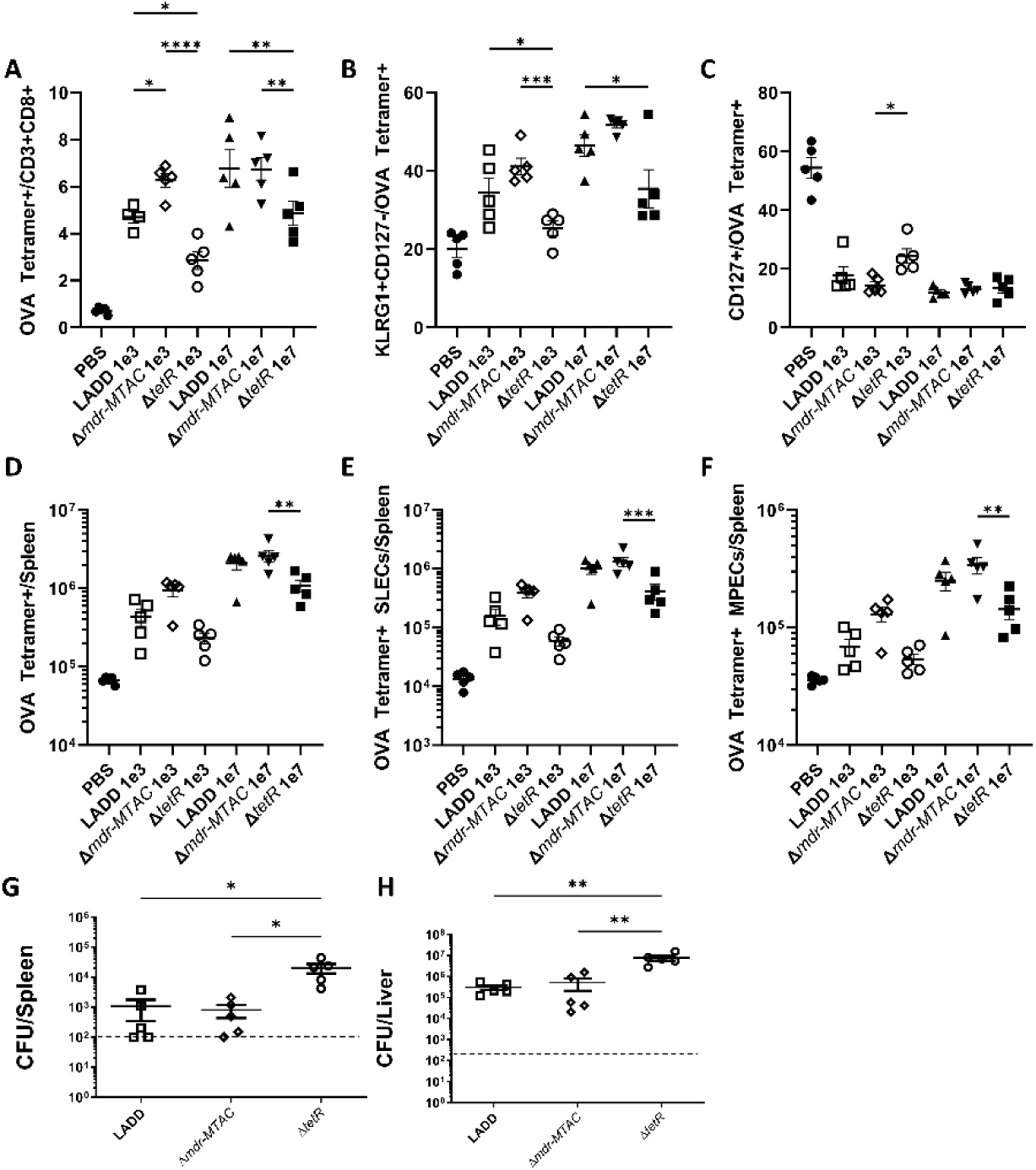
The T-cell response is enhanced in mice immunized with type I Interferon hypo-inducing *L. monocytogenes,* but protective immunity remains unchanged. Mice were immunized with indicated strain and dose of *L. monocytogenes* and OVA-specific T-cell A) frequencies and D) numbers per spleen, or OVA-specific SLEC B) frequencies and D) numbers per spleen, or OVA-specific MPEC C) frequencies and F) numbers per spleen, were assessed. Mice were immunized with 1×10^3^ of the indicated strain expressing OVA and B8R and challenged with 5×10^5^ WT *L. monocytogenes* expressing OVA and B8R one month later. Three days post challenge, livers and spleens were harvested and plated for CFU in G) spleens and H) livers. Data shown are representative of two independent experiments of 4-5 mice per group. Significance was determined by a one-way ANOVA with Tukey’s correction for panels A-F or a Kruskal-Wallace non-parametric test with Dunn’s correction for panels G, H. * p < 0.05.

Δ*mdr-MTAC* vaccination improved T-cell responses compared to WT, thus we hypothesized that altering type I IFN during the primary T-cell response would improve protective immunity to rechallenge. To test this hypothesis, we immunized mice with a low dose (10^3^) of the indicated strain expressing B8R and OVA and rechallenged them 30 days later with 5 LD50s of WT *L. monocytogenes* expressing B8R and OVA and assessed CFU in the livers and spleens 3 days post-challenge. In contrast to the enhanced T-cell response observed for *Δmdr-MTAC*, we saw no enhanced protection from rechallenge compared to WT in either organ (Figure 5G-H). In line with previous results^14^, we observed that *ΔtetR* reduced protection. Therefore, although we observe enhanced SLEC and MPEC frequencies during the primary T-cell response, this did not translate to enhanced protection and may suggest that type I IFN in the secondary response from a WT strain compared to *Δmdr-MTAC* used during immunization can blunt enhancements observed in the primary T-cell response.

### Antigen Presenting Cells Are Sensitive to type I IFN Signaling

Type I IFN plays a major role in T-cell priming and thus the memory pool. We initially hypothesized that type I IFN affects T-cells directly, however, Archer et al (2014) recently demonstrated the impact of type I interferon on T-cell responses is T-cell extrinsic by adoptively transferring WT or IFNAR^−/−^ T-cells followed by *L. monocytogenes* vaccination^14^. We, therefore, hypothesized that type I IFN impairs T-cell responses through signaling on antigen-presenting cells (APCs). To test this hypothesis, we performed *ex vivo* T-cell proliferation assays using APCs from WT or IFNAR^−/−^ mice, from B16-Flt3L tumor-bearing WT or IFNAR^−/−^ mice as previously described^33^. DCs were then infected with our vaccine strain expressing B8R and OVA and after 6 hours of infection, we added CFSE-labeled OT-I T-cells. At varying time points after co-culture, we assayed for proliferation by CFSE dilution (Fig. S10C) and for IFNγ production in the supernatants by ELISA. As a control, we loaded DCs with SIINFEKL peptide in the absence of *L. monocytogenes* infection to promote T-cell proliferation without type I IFN induction. While SIINFEKL-loaded WT and IFNAR^−/−^ DCs induced comparable robust T-cell proliferation, only IFNAR^−/−^ APCs induced robust T-cell proliferation comparable to SIINFEKL-loaded APCs when antigen was supplied by *L. monocytogenes* infection (Figure 6A). In fact, we speculate that *L. monocytogenes* may be inducing a suppressive factor since IFNAR^−/−^ APCs induced the same proliferation (80-90% of T-cells dividing at least once) as SIINFECKL loaded APCs, while WT was reduced to half that. Alterations in T-cell priming were observed as early as day two post-co-culture, and no proliferation was observed on day one (Fig. S10 A-B). In addition to inducing increased T-cell proliferation, IFNAR^−/−^ APCs induced greater IFNγ production by at least two-fold (Figure 5B). Collectively, these data suggest that IFNAR signaling on APCs inhibits optimal T-cell priming in the context of *L. monocytogenes* immunization.

**Figure 6 -.**
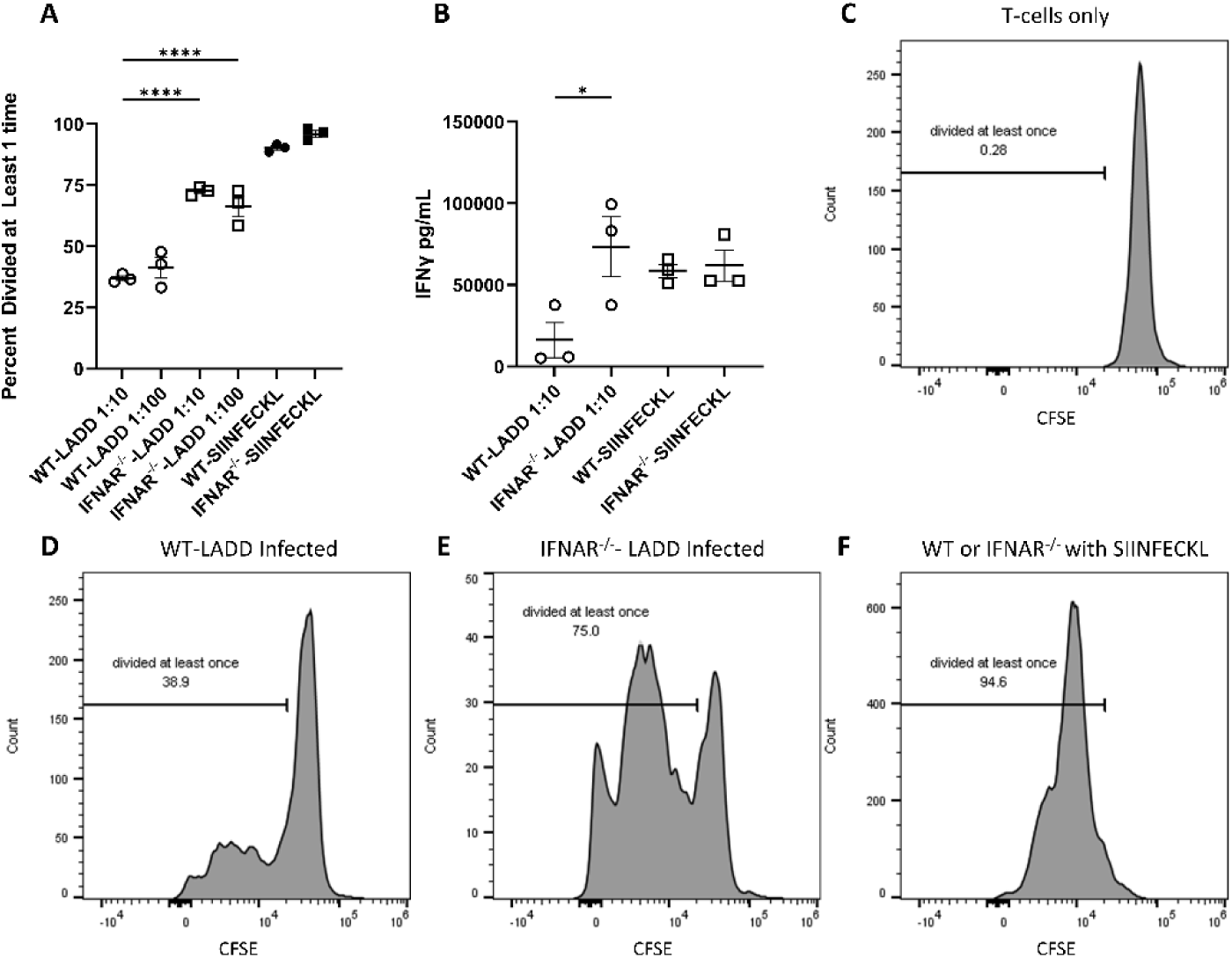
APCs are responsive to type I IFN during *ex vivo* T-cell co-culture after infection with *L. monocytogenes.* CD11c+ splenocytes were isolated from B16-Flt3L tumor-bearing WT or IFNAR^−/−^ mice and infected with *L. monocytogenes* expressing OVA and B8R for 6 hours or loaded with 2μM SIINFEKL peptide for two hours. Then CFSE-labeled OT-I T-cells were added at varying T-cell:APC ratios. The percentage of proliferated CD3+CD8+ T-cells was determined by flow cytometry for CFSE dilution at A) 72 hours and B) IFNγ was assessed by ELISA. Representative flow cytometry histograms showing T-cell proliferation are presented for C) no APCs, D) WT APCs, E) IFNAR-^1^_-_ APCs, and F) SIINFEKL-Loaded APCs. Data shown are representative of two independent experiments. Significance was determined by a one-way ANOVA with Bonferroni’s correction. * p < 0.05.

## Discussion

L. monocytogenes is a potent activator of CD8^+^ T-cells and has been used as a tool to study cell-mediated immunity and as a vaccine platform in the context of tumor immunotherapy. Optimized *L. monocytogenes* vaccine development demands a better understanding of how *L. monocytogenes* activates robust cell-mediated immunity and the parameters that impact the quantity and quality of this response. In this study, we demonstrated that type I IFN impairs T-cell responses, specifically by blunting MPEC formation leading to deficits in the memory pool and subsequent recall response that ultimately compromise protective immunity. By blocking this pathway, we enhanced all these metrics. We examined the role of inflammasome activation and found its impact is minimal, likely due to the low level of inflammasome activation by *L. monocytogenes*. Utilizing vaccine strains genetically engineered to induce less type I IFN, we overcame deficits in T-cell immunity caused by type I IFN, although there is still room for improvement. Finally, we show the detrimental effect of type I IFN is at least partially through IFNAR signaling on the antigen-presenting cells rather than on T-cells as was previously hypothesized. In total, we reveal that type I IFN is a rational pathway to target for intervention during vaccine design, presented one such strategy to overcome deficits resulting from type I IFN signaling, and highlighted a specific cell subset to target in other therapeutic interventions involving type I IFN signaling. These findings justify further studying the impact of innate immune signaling pathways on T-cell responses in a cell-specific manner.

We were surprised that Caspase-1^−/−^ mice had comparable T-cell responses to WT. However, *L. monocytogenes* is quite adept at avoiding the inflammasome^12^. Thus we speculate that due to the low level activated by *L. monocytogenes,* inflammasome activation has a less crucial role than was previously hypothesized^13^. Our results contrast with Rana et. al (2020) who observed that Caspase-1 was necessary for optimal T-cell responses to *L. monocytogenes*^34^. Our strain is deficient for the host-actin assembly protein ActA to attenuate its virulence and is cleared in 2-3 days^20^ while Rana et al. (2020) utilized WT *L. monocytogenes* which persisted for nearly 7 days-post immunization and is not suitable as a vaccine strain. However, using a strain of *L. monocytogenes* that hyperactivates the inflammasome, we demonstrated that this is detrimental to the T-cell response^13^, suggesting inflammasome activation should not be pursued for a more efficacious vaccine. We did uncover a role for inflammasome activation in the absence of type I IFN signaling as Caspase-1^−/−^/IFNAR^−/−^ mice had worse T-cell immunity than IFNAR^−/−^ alone. This may suggest a common signaling nodule exists between the two pathways. In the context of *Mycobacterium tuberculosis* infection, a PGE_2_, IL-1, and type I IFN signaling axis regulates protective immunity such that IL-1 signaling was necessary to promote PGE_2_ production while type-I IFN signaling blunted PGE_2._ production^35^. We recently demonstrated PGE_2_ is necessary for optimal T-cell responses toward *L. monocytogenes*^36^. Thus, we hypothesize that type I IFN impairs T-cell responses, in part, through diminishing PGE_2_ production. Repression would be relieved in IFNAR^−/−^ mice to promote enhanced T-cell responses, however, because PGE_2_ production may be dependent on IL-1 signaling, the benefit of IFNAR blockade may be lost by additionally blocking inflammasomes. Whether type I IFN impairs PGE_2_ production in the context of *L. monocytogenes* infection, or if inflammasome activation is necessary for PGE_2_ production remains to be determined. Future efforts on this topic are ongoing and may lead to further optimization of vaccine design through modulating PGE_2_ production.

Multiple possibilities exist for why type I IFN signaling on APCs is detrimental to T-cell immunity during *L. monocytogenes* vaccination. IFNAR^−/−^ APCs may be more phagocytic, more permissive to *L. monocytogenes* replication, better at antigen presentation, have enhanced co-stimulatory receptor expression or reduced co-inhibitory receptor expression, or provide a more optimal inflammatory milieu (for example more PGE_2_). Recent studies suggest higher type I IFN expression is correlated with increases in expression of the apoptosis-inducing ligand, TRAIL, and its receptor, DR5, in bulk lymphoid tissue in human HIV patients^39^. Thus, TRAIL expression induced specifically on APCs during *L. monocytogenes* vaccination may promote antigen-specific T-cell apoptosis during T-cell/APC interactions. Consistent with this idea, macrophage-produced IFN-β leads to increased TRAIL expression on myeloid cells during influenza A infection^37^. And like the phenotype observed for IFNAR^−/−^ mice^38^, TRAIL deficient mice are more resistant to *L. monocytogenes* infection and show reduced cell death during early *L. monocytogenes* infection^39^. However, the antigen-specific T-cell response was not explored in this study. In line with this hypothesis, type I IFN signaling promotes apoptosis in germinal centers during *L. monocytogenes* infection^40^ and may result in apoptotic body formation, which ultimately leads to IL-10 production^12^. Supporting this notion, IFNAR^−/−^ mice produce less IL-10 than WT during *L. monocytogenes* infection^41^ and IL-10 deficient mice have enhanced T-cell responses to *L. monocytogenes*^42,43^. Future experiments should test if deficits in T-cell responses from *ΔtetR* are rescued in an IL-10 deficient mouse and interrogate immunosuppressive and/or cell death ligands on APCs and on interacting cell subsets during *L. monocytogenes* vaccination.

To improve *L. monocytogenes-*induced T-cell immunity, several approaches could be used. While type I IFN blockade is reasonable for generating more antigen-specific T-cells during vaccination, type I IFN signaling is critical for the generation of anti-tumor immunity in tumor-bearing mice^44^. Thus, temporally blocking type I IFN only during vaccination may result in enhanced anti-tumor immunity. Similarly, the role of type I IFN for other non-vaccine strategies aimed at generating anti-tumor T-cells such as in situ vaccines or radiotherapy has not been thoroughly dissected^45^ and warrants investigation. Temporal blockade could be achieved using interferon blocking antibodies such as Sifalimumab which is already approved for the treatment of Lupus and poliomyelitis or Jak/STAT inhibitors. Alternatively, vaccine strains that further reduce CDN secretion or secrete STING antagonists to dampen STING-dependent type I IFN production may be another rational strategy. In a similar vein, if IL-10 proves to be the critical downstream mediator of type I IFN-induced deficits in T-cell immunity, IL-10 blocking antibodies could be deployed during vaccination, which would bypass the need to block type I IFN and potentially spare any beneficial impacts type I IFN has in generating anti-tumor immunity. Finally, other innate immune pathways could be modulated to enhance the *L. monocytogenes-*driven T-cell response. As alluded to earlier, we recently discovered production of PGE_2_, a lipid mediator of inflammation, is critical for optimal T-cell responses to *L. monocytogenes*^21^. *L. monocytogenes* could be engineered to express the biosynthetic pathway for PGE_2_ synthesis or secrete the metabolic products necessary to enhance PGE_2_ production. Current efforts are underway to determine how cytosolic *L. monocytogenes* promotes PGE_2_ production. Another important avenue to explore will be how PGE_2_ impacts memory precursor formation and maintenance. If such a link exists, we predict loss of PGE_2_ should result in decreased MPECs and would suggest another route to improve vaccination strategies as enzymatic inhibitors of PGE_2_ production (ibuprofen) are often given during vaccination to limit side effects.

In total, our work demonstrates that *L. monocytogenes* vaccines can be improved by blocking type I IFN, the type I IFN responsive cells are APCs, and that there is room for more improvement through examination of crosstalk with alternative pathways, known and unknown, activated by *L. monocytogenes*.

## Materials and Methods

### Ethics Statement

This work was carried out in strict accordance with the recommendations in the Guide for the Care and Use of Laboratory Animals of the National Institutes of Health. All protocols were reviewed and approved by the University of Wisconsin-Madison Institutional Animal Care and Use Committee.

### Mouse Strains

6-16 week C57BL/6 female mice were obtained from the NCI and Charles River NCI facility and were age matched to their experimental cohort when possible. Caspase-1^−/−^ (Casp1^tm1Flv^) and IFNAR1^−/−^ (B6.129S2-Ifnar1^tm1Agt/Mmjax^) mice were obtained from the Jaxson Laboratory and were maintained as homozygous breeding pairs and were periodically genotyped. OT-I (C57BL/6^Tg(TcraTcrb)1100Mjb/J^) mice were a kind gift from Dr. Susana Fabry and maintained at homozygosity. Caspase-1^−/−^;IFNAR^−/−^ were generated by crossing Caspase-1^−/−^ and IFNAR^−/−^ homozygous mice to generate heterozygotes and incrossing those offspring to generate Caspase-1^−/−^;IFNAR^−/−^. From thereon, mice were maintained at homozygosity.

### Bacterial strains, cloning, and growth conditions

*L. monocytogenes* (strain 10403S) was routinely grown at 30°C for overnight cultures or 37°C for back-diluted cultures in brain heart infusion (BHI) broth (BD, 237500) or on BHI + 1.5% agar (Fisher, BP1423500) plates and frozen in BHI + 40% glycerol (Acros Organics, 332030025). *E. coli* was routinely grown at 37°C in LB or on LB + 1.5% agar plates and frozen in LB + 40% glycerol. SM10 was used for conjugating plasmids into *L. monocytogenes*. Antibiotics were used at the following concentrations: 200 μg/ml streptomycin (Fisher, BP910-50),10 μg/ml chloramphenicol (Dot Scientific, DSC61000-25), 2 μg/ml erythromycin (Dot Scientific, DSE57000-10). The attenuated (LADD) strain used in the analysis of T-cell responses was in the Δ*actA*Δ*inlB* background as previously described and engineered to express full-length OVA and the B8R20-27 epitope^20^. Phage transduction in *L. monocytogenes* was performed as previously described^46^ using U153 phage make strain LADD-Δ*tetR-*B8R-OVA from LADD-B8R-OVA and Δ*tetR*^32^. Strain LADD-Δ*mdrMTAC-*B8R-OVA was constructed by conjugating plasmid pPL2-ActAN100-B8R-OVA into LADD-Δ*mdrMTAC* (a kind gift from Dr. Anat Herskovits)^31^ using SM10 *E. coli*.

### *In vivo* immunizations

The indicated strains were grown in BHI at 30°C overnight without agitation. Cultures were then back diluted 1:5 in BHI and grown at 37°C with agitation until log phage (OD 0.4-0.6, ~1-1.5 hours). Bacteria were diluted in PBS and mice were infected with the indicated dose intravenously in 200 µL of PBS solution (Fisher, 14190250). For T-cell cell analysis, spleens were harvested at the indicated timepoints post-immunization. For challenge experiments, mice were immunized with the indicated strain and dose. 30-33 days post immunization mice were challenged with 5×10^5^ WT *L. monocytogenes* expressing B8R and OVA. Three days post challenge, spleens and livers were collected and homogenized in 0.1% IPEGAL (Sigma, I8896- 50ML) in PBS and plated on BHI plates containing streptomycin to quantify bacterial loads.

### Flow Cytometry and T-cell *ex vivo* Stimulations

Mice were sacrificed at the indicated time points, and spleens were collected and smashed through 70 µm mesh screens (Falcon, #352350) as previously described^11^. Briefly, cells were lysed in ACK lysis buffer (0.15M NH_4_Cl [Sigma, A9434-500G], 10mM KHCO_3_ [Fisher, 298-14-6], 0.1mM Na_2_EDTA [Sigma, E7889]) for less than 10 minutes and resuspended in T-cell Media (RPMI [Fisher, #11875093], with 10% FBS [Fisher, SH30071], 1% L-Glutamine [Fisher, #25030081], 1% Sodium Pyruvate [Invitrogen, #11360070], 0.1% β-mercaptoethanol [Fisher, #21985023], 1% HEPES [Invitrogen, 15630-080], 1% MEM Non-essential amino acids [Invitrogen, #11140-050], 1% Antibiotic/Antimycotic [Fisher, 15240062]) and counted using a Z1 Coulter counter. For tetramer staining, 2×10^6^ cells were immediately blocked for Fc (Fisher, #501054562) at 1:100 dilution in FACs buffer (1% heat inactivated FBS in PBS) for 30 minutes on ice, then stained using OVA or B8R tetramer at 1:300 (Bv421, NIH Tetramer Facility, Atlanta, GA) for 30 minutes on ice. Cells were washed twice in FACs and then stained for extracellular markers CD3 (PE/Cy7, 1:50, clone 145-2C11), CD8α (FITC, 1:200, clone 53-6.7), B220 (AF700, 1:160, clone RA3-6B2), CD127 (APC, 1:50, clone A7R34), and KLRG1 (PE, 1:200, clone 2F1) for 30 minutes on ice. Cells were then fixed in IC Fixation buffer (Invitrogen, #FB001) for 20 minutes on ice. The reaction was stopped by the addition of an equal volume of FACs, and cells were stored overnight prior to flow.

For peptide stimulation, 2×10^6^ splenocytes were plated in 96-well plates (Fisher, #130188) in 200 μL T-cell media and stimulated with the indicated peptide (SIINFEKL [Anaspec, #AS-60193-1], or TSYKFESV [Anaspec, #AS-64688]) at a final concentration of 2 µM for 5 hours prior to Fc block as before, then surface staining with CD3 (PE/Cy7, 1:50) and CD8α (FITC, 1:200, clone 53-6.7). Cells were then fixed as before, and subjected to permeabilization buffer (Fisher, #00-8333-56) followed by staining in permeabilization buffer with IFNγ (APC, 1:300, clone XMG1.2), TNFα (PerCP/Eflour710, 1:500, clone MP6-XT22), and IL-2 (PE, 1:120, clone JES6-5H4). Cells were then washed in FACs twice and stored overnight prior to acquisition using an LSRII flow cytometer (BD Biosciences, San Jose, CA) or an Attune NxT flow cytometer (Thermo Fisher Scientific) with manufacturer-provided acquisition software and analyzed with FlowJo software (Tree Star, Ashland, OR).

### BMDM generation and culture

Bone marrow-derived macrophages (BMDMs) were made using six-to eight-week-old C57BL/6 mice or indicated strains as previously described^13^. Briefly, bone marrow was harvested from the long bones of mice and cultured in the presence of M-CSF from transfected 3T3 cell supernatant for 6 days with a supplement of M-CSF at day 3 and frozen down in media + 10% DMSO + an additional 10%FBS until use. BMDM generation media was RPMI (Invitrogen, #11875119) + 20% FBS, 10% M-CSF (from 3T3 cells), 1% L-Glutamine (Fisher, #25030081), 1% Sodium Pyruvate (Invitrogen, #11360070), 0.1% β-mercaptoethanol (fisher, #21985023). 5×10^5^ BMDMs were cultured in BMDM culture media (the same as BMDM generation media but with 10% FBS only) and plated in a 24 well dish (Fisher, #12556006) in 0.5 mL BMDM media, and allowed to adhere for 18 hours and infected at the indicated MOI with the indicated strain grown overnight at 30°C without agitation. After 1 hour, gentamycin (VWR, #12001-684) was added to a final concentration of 50µg/mL to kill extracellular bacteria. 6 hours after infection, the plate was spun, and the media was removed and stored at 4°C until use in ELISA.

### Cytokine analysis

IFN-β ELISA was performed as previously described^22,47–49^ with slight modifications. Half-volume ELISA plates (Fisher, #07-200-37) were coated overnight with 100 µL rat-α-mouse IFN-β antibody (Santa Cruz, # sc-57201) at 0.1 µg/mL in plate coating buffer (0.1 M carbonate [Thermo, 25080-094], pH 9) at 4°C. The solution was removed, and the plate was washed with 200 µL wash solution (1% FBS, 0.05% Tween [Sigma, P9416-100ML], in PBS). The solution was removed, and the plate was blocked with 200 µL blocking solution (1% BSA [Sigma, #A9418-5G], 0.05% Tween [Sigma, #P9416-100ML], in PBS). The solution was removed and sample or standard curve (made with mouse IFN-β [PBL Biomedical Laboratories, #12400-1] in blocking buffer) was added to wells and incubated overnight at 4°C. The solution was removed, and plate was washed 2X with 200 µL wash buffer. 100 µL rabbit-α-mouse IFN-β polyclonal antibody (PBL assay science, #32400-1) at 1:2000 dilution in blocking buffer was added and the plate was incubated for 2 hours at room temperature. The solution was removed and the plate was washed 2X with 200 µL wash solution then 100 µL goat-α-rabbit HRP (cell signaling, #7074S) at 1:2000 in blocking buffer was added. After 1 hour incubation, the solution was removed and the plate was washed 2X with 200 µL wash solution and 100 μL TMB solution (Sigma, #T0440-100ML) is added. The plate was incubated for 30 minutes before the addition of 100 µL 1 M H_2_SO_4_. Absorbance was read at 450 nm in an Eon Microplate Spectrophotometer (BioTekInstruments, Inc., Winooski, VT).

IL-1β ELISA was performed according to manufactures instructions using the Invitrogen IL-1 beta Mouse Uncoated ELISA Kit (Invitrogen, #88701388) with the exception that half-volume ELISA plates were used.

### Ex vivo T-cell proliferation Assay

B16 cells expressing FMS-like tyrosine kinase 3 ligand (Flt3L) were cultured in RP10G Media (RPMI [Fisher, #11875093], with 10% FBS, 1% L-Glutamine [Fisher, #25030081], 1% Sodium Pyruvate [Invitrogen, #11360070], 0.1% β-mercaptoethanol [Fisher, #21985023], 1% HEPES [Invitrogen, 15630-080], and 0.05 mg/mL gentamycin [Invitrogen, #15750-060]). 2×10^6^ B16-FLT3L cells implanted into the rear right flank of WT or IFNAR^−/−^ mice. Two to three weeks post-implantation, when tumors had reached approximately 0.5-1 cm in greatest dimension, spleens were removed and mashed through 70 μm mesh strainers in ACK buffer and resuspended in EasySep Buffer (StemCell, #20144). Cells were counted using a Z1 Coulter counter and CD11C+ cells were isolated using the EasySep Mouse Pan-DC Enrichment Kit (StemCell, 19763) according to manufacturer’s instructions. Cells were routinely 70-95% CD11C-positive (CD11c FitC, biolegend, 117306, 1:200), and CD86-negative (CD86 APC, Ebiosciences, 17-0862-81, 1:500) prior to plating as assessed by flow cytometry. After isolation, cells were spun at 500xG and resuspended in Flt3L DC Buffer (T-cell buffer + 10% B16-Flt3L conditioned RP10G medium + 1000U/mL Mouse GM-CSF [BD Biosciences, #554586]). Dendritic cells were plated in a total volume of 75 μL and rested overnight. The next morning, cells were infected with the indicated strain in 5 μL Flt3L DC Buffer at MOI = 2 for one hour prior to the addition of gentamycin (50 μg/mL). At 4 hours post infection, chloramphenicol is added to a final concentration of 10μg/mL to cease intracellular replication of *L. monocytogenes*. In separate conditions, OVA peptide is added for two hours to a final concentration of 2μM. During infection OT-I T-cells were isolated from the spleens of homozygous mice using the EasySep™ Mouse CD8^+^ T Cell Isolation Kit (StemCell, 19853). CD8^+^ T-cells were routinely 95-99% pure as assessed by flow cytometry and subsequently labeled with CFSE. CSFE labeling was accomplished by incubating isolated T-cells in T-cell media + 10μM CFSE (StemCell, #75003) for 10 minutes prior to 3 washes in T-cell buffer. At six hours post-infection or two hours post-OVA peptide addition, 100,000 CFSE (StemCell, 75003) labeled OT-I T-cells are added in Flt3L DC Buffer to bring the final volume to 100 μL. At the indicated timepoints, media was collected and frozen at −80°C and cells were labeled with Ghost Dye Red 780 (Tonbo, #13-0865-T500) according to manufacturer’s instructions prior to staining for CD8-BV421 as above and acquired using an LSRII flow cytometer (BD Biosciences, San Jose, CA) or an Attune NxT flow cytometer (Thermo Fisher Scientific) with manufacturer-provided acquisition software and analyzed with FlowJo software (Tree Star, Ashland, OR). The percentage of divided or undivided cells was then calculated by comparison to T-cells cultured without APCs.

### Statistics and Analysis

Statistical analysis was performed by GraphPad Prism Software (La Jolla, CA) and analyzed via Kruskal-Wallace, Mann Whitney or one-way ANOVA with Tukey’s correction as indicated.

## Acknowledgements

This work was funded by the National Institutes of Health R01CA188034 (JDS). Additionally, this material is based upon work supported by the National Science Foundation Graduate Research Fellowship Program (ZTM) under Grant No. DGE-1747503. Any opinions, findings, and conclusions or recommendations expressed in this material are those of the author(s) and do not necessarily reflect the views of the National Science Foundation. We would also like to thank the NIH Tetramer Core Facility for provision of MHC-I B8R tetramers. The authors thank the University of Wisconsin Carbone Cancer Center Flow Cytometry Laboratory, supported by P30 CA014520, for use of its facilities and services.

---

**Supplemental Figure 1 –.**
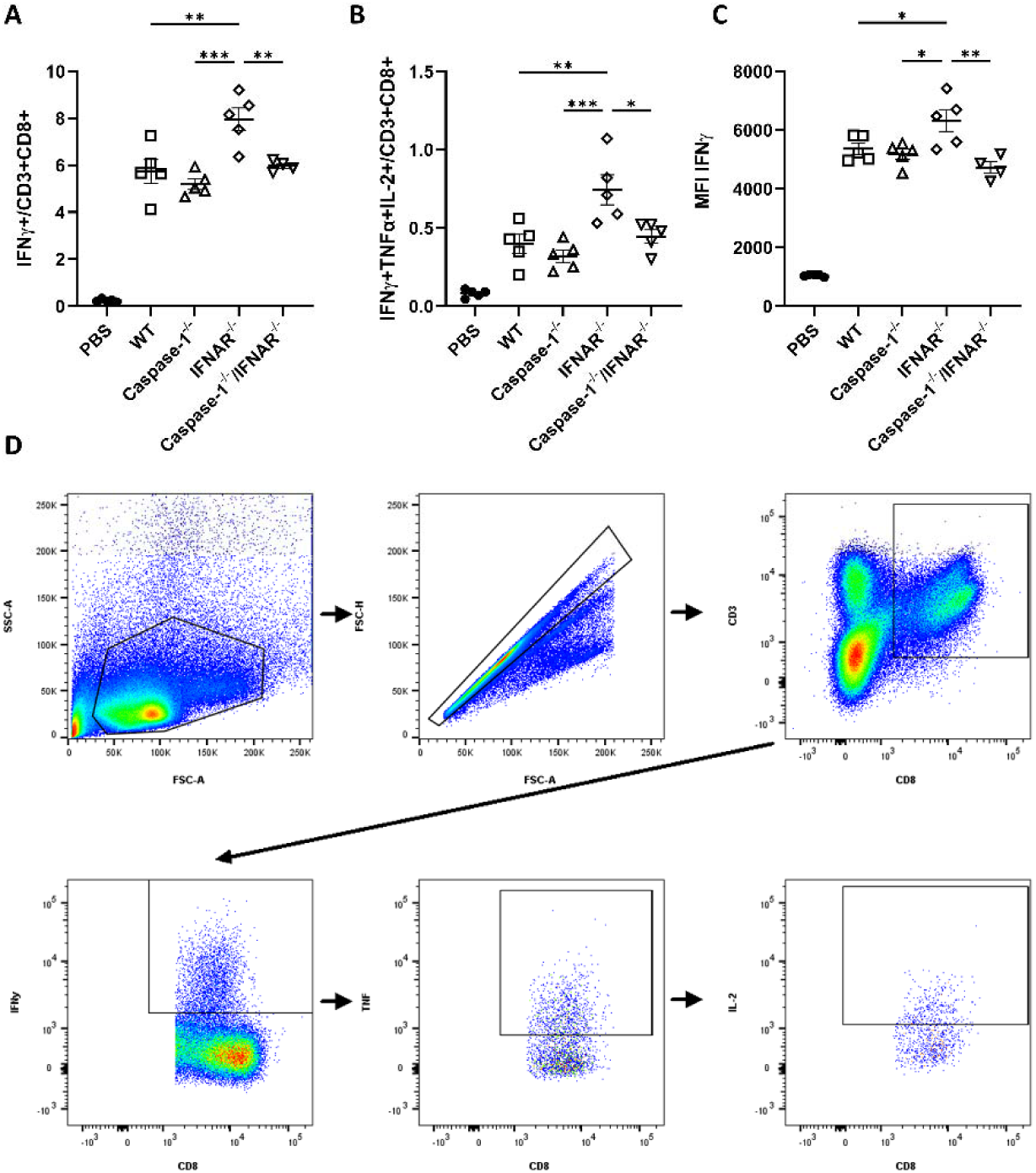
B8R-specific antigen responses in T-cell response and gating strategy for peptide stimulated splenocytes (goes with figure 1). Mice were immunized with 10^3^ LADD expressing B8R and OVA. Splenocytes were examined for B8R specific antigen responses by *ex vivo* peptide stimulation 7 days after immunization. Cells were gated for CD3+ and CD8+ followed by A) IFNγ or B) IFNγ+ TNFα+ and IL-2+ as well as C) IFNγ expression by MFI. Data are representative of two independent experiments of 4-5 mice per group Significance was determined by a one-way ANOVA with Tukey’s correction. * p < 0.05. D) Cells were gated on 1) Lymphocytes, 2) single cells, 3) CD3+CD8+, 4) IFNγ+, 5) TNFα+, 6) IL-2+. Gates were drawn based on FMOs.

**Supplemental figure 2 -.**
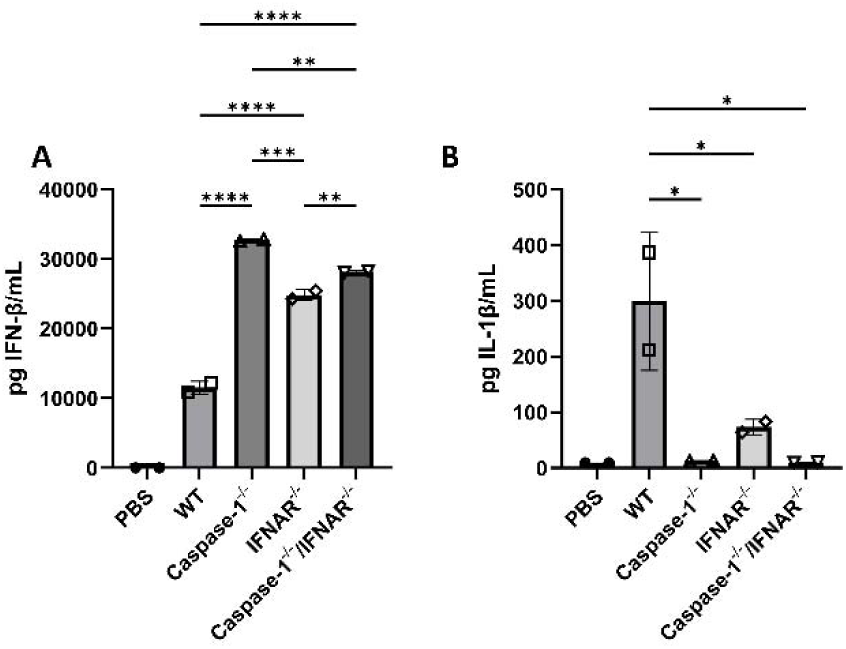
BMDMs produce IFN-β and IL-1β after infection with *L. monocytogenes e*(goes with figure 1). Wild-type, Caspase-1^−/−^, IFNAR^−/−^ or Caspase-1^−/−^/IFNAR^−/−^ BMDMs were infected with attenuated *L. monocytogenes* at an MOI = 5 +/− PAM3CSK4 and assessed for A) IFN-β or B) IL-1β production in the supernatants by ELISA at 6 hours post infection. Data are representative of two independent experiments. Significance was determined by one-way ANOVA with Tukey’s correction. *p<0.5.

**Supplemental Figure 3 -.**
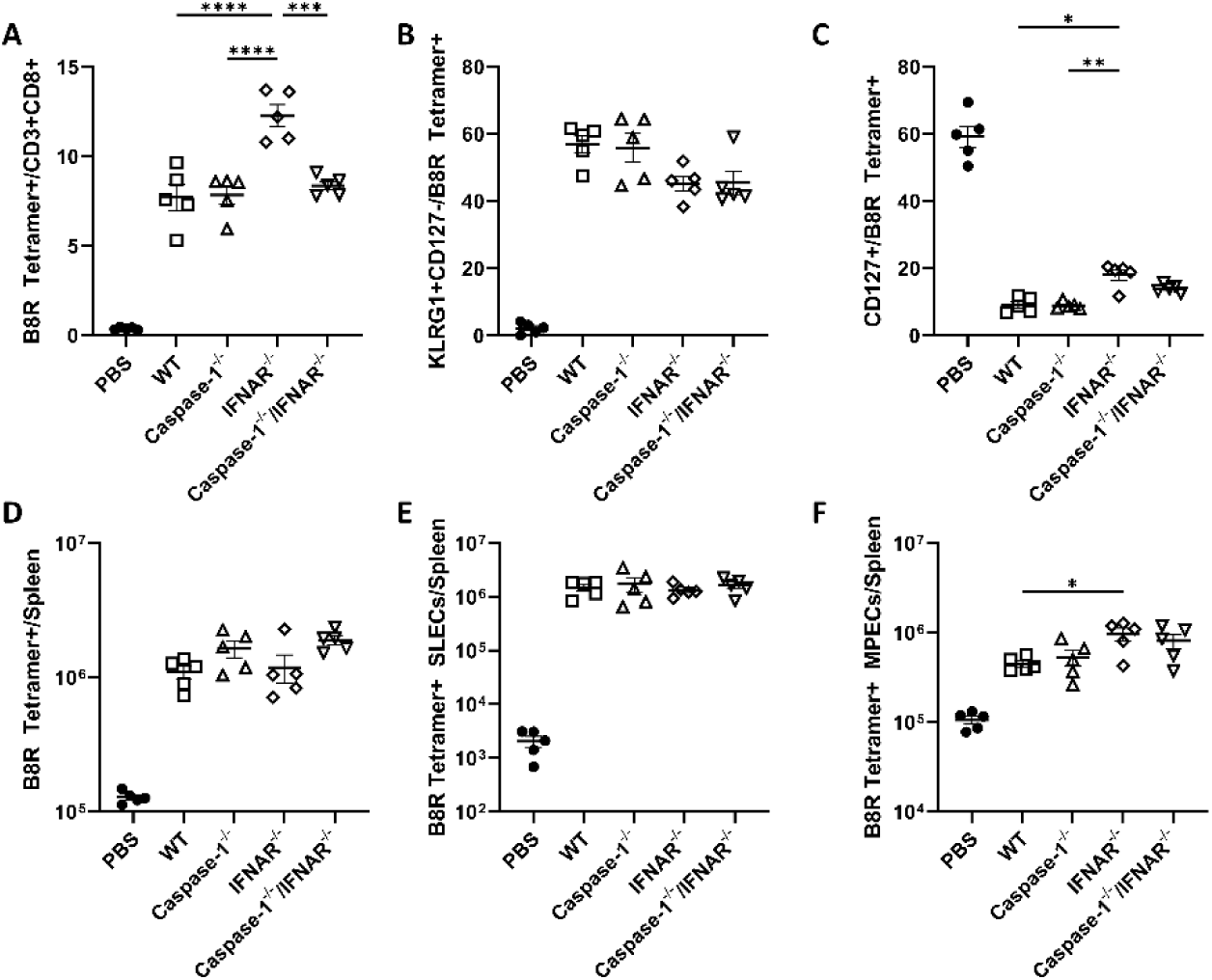
B8R specific SLEC and MPEC analysis at primary T-cell response (goes with figure 2) Mice were immunized with10^3^ cfu *L. monocytogenes* expressing B8R and OVA. B8R-specific responses were examined at day 7 post immunization by gating for CD3+ CD8+ and B220-cells and examing A, D) B8R-tetramer+ or B, E) B8R-tetramer+ and KLRG1+ (SLECs) or C, F) B8R-tetramer+CD127+ cells (MPECs). Data shown are representative of two independent experiments of 4-5 mice per group. Significance was determined by a one-way ANOVA with Tukey’s correction. * p < 0.05.

**Supplemental Figure 4 (goes with figure 2) –.**
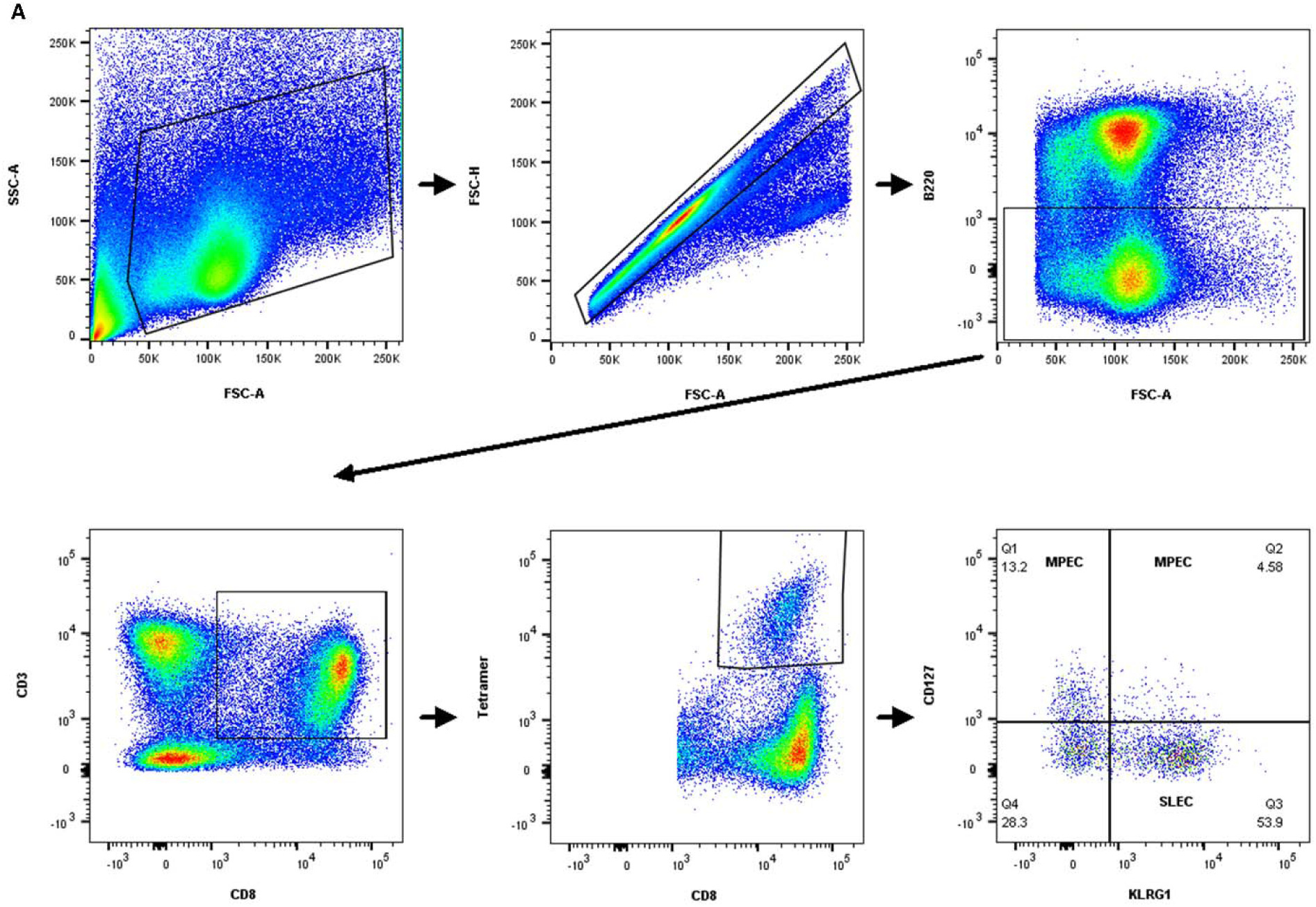
Gating strategy for tetramer panels. Cells were gated on 1) Lymphocytes, 2) single cells, 3) B220 negative (to remove sticky B-cells), 4) CD3+CD8+, 5) Tetramer+, 6) CD127 and KLRG1. Gates are drawn according to FMOs.

**Supplemental Figure 5 -.**
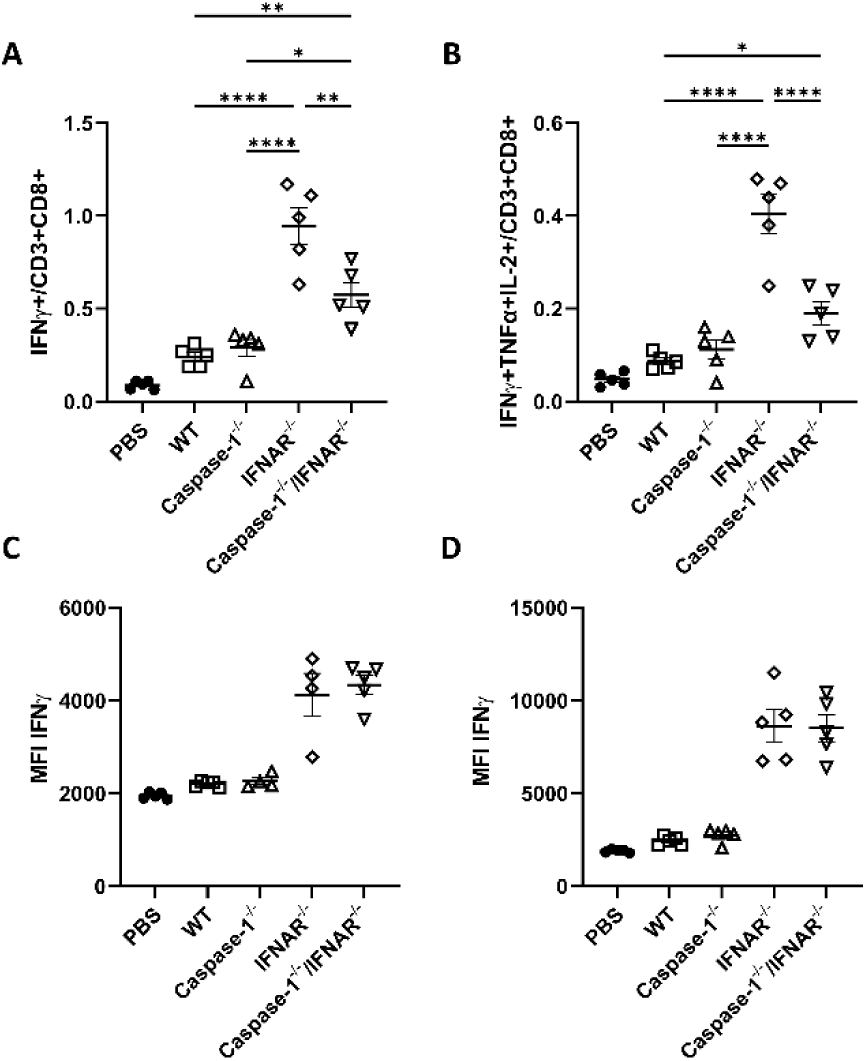
B8R-specific antigen responses in T-cell response and IFNγ expression for peptide stimulated splenocytes at memory phase (goes with figure 3). Mice were immunized with 10^3^ LADD expressing B8R and OVA. Splenocytes were examined for B8R specific antigen responses by *ex vivo* peptide stimulation 30-32 days after immunization. Cells were gated for CD3+ and CD8+ followed by A) IFNγ or B) IFNγ+ TNFα+ and IL-2+ as well as for IFNγ expression by MFI for C) OVA-specific or D) B8R-specific T-cells. Data are representative of two independent experiments of 4-5 mice per group Significance was determined by a one-way ANOVA with Tukey’s correction. * p < 0.05.

**Supplemental Figure 6 -.**
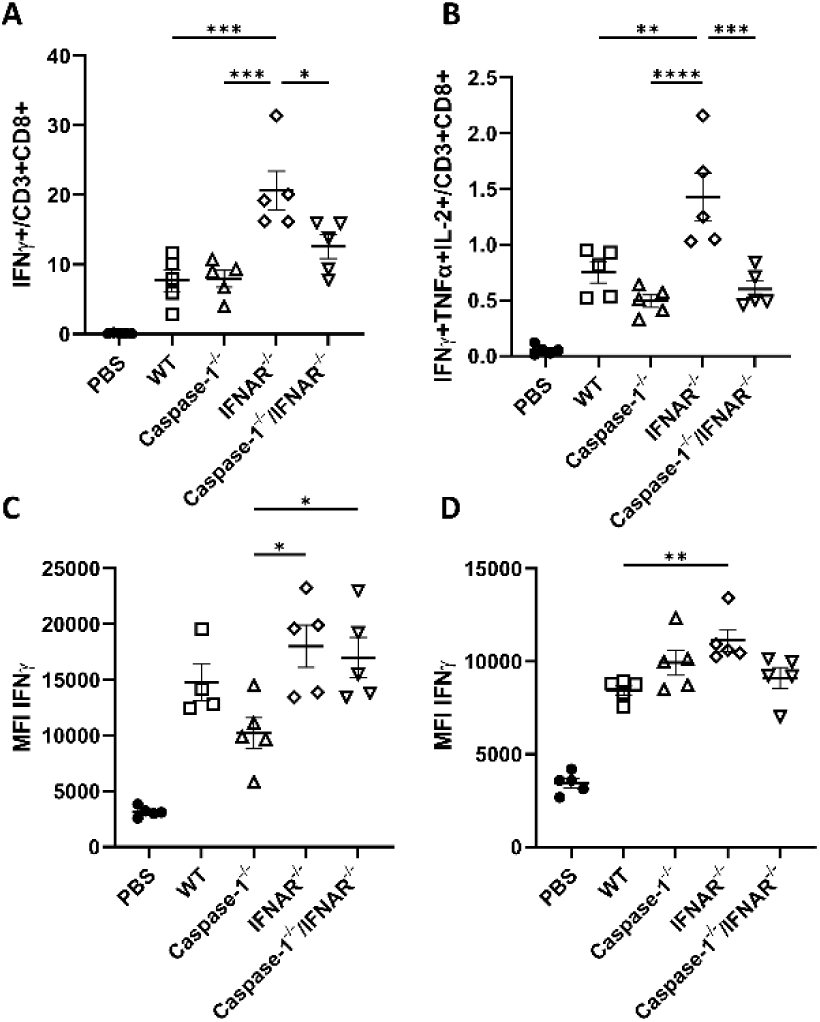
B8R-specific T-cell responses and IFNγ expression for peptide stimulated splenocytes during recall response (goes with figure 4). Mice were immunized with 1×10^3^ *L. monocytogenes* expressing B8R and OVA and then challenged with 5×10^5^ WT *L. monocytogenes* expressing B8R and OVA. B8R-specific responses were examined by peptide stimulation at day 3 post challenge for CD8+ T-cells producing A) IFNγ, and B) IFNγ+TNFα and IL-2+, as well as IFNγ expression by MFI for C) OVA-specific or D) B8R-specific T-cells. Data are representative of two independent experiments of 4-5 mice per group Significance was determined by a one-way ANOVA with Tukey’s correction. * p < 0.05.

**Supplemental Figure 7 (goes with figure 5) -.**
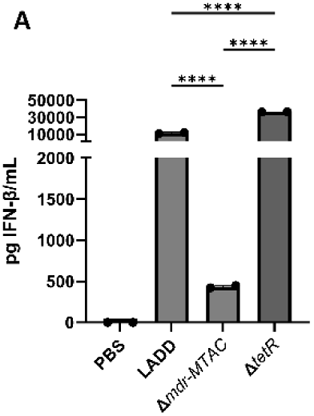
*Listeria* engineered to modify CDN secretion machinery induce altered levels type I IFN production. Bone marrow-derived macrophages were infected with the indicated strain at an MOI = 5, after 30 minutes, media was replaced with gentamicin-containing media. Infection proceeded for 6 hours before performing ELISA for IFN-β on supernatants.

**Supplemental Figure 8 –.**
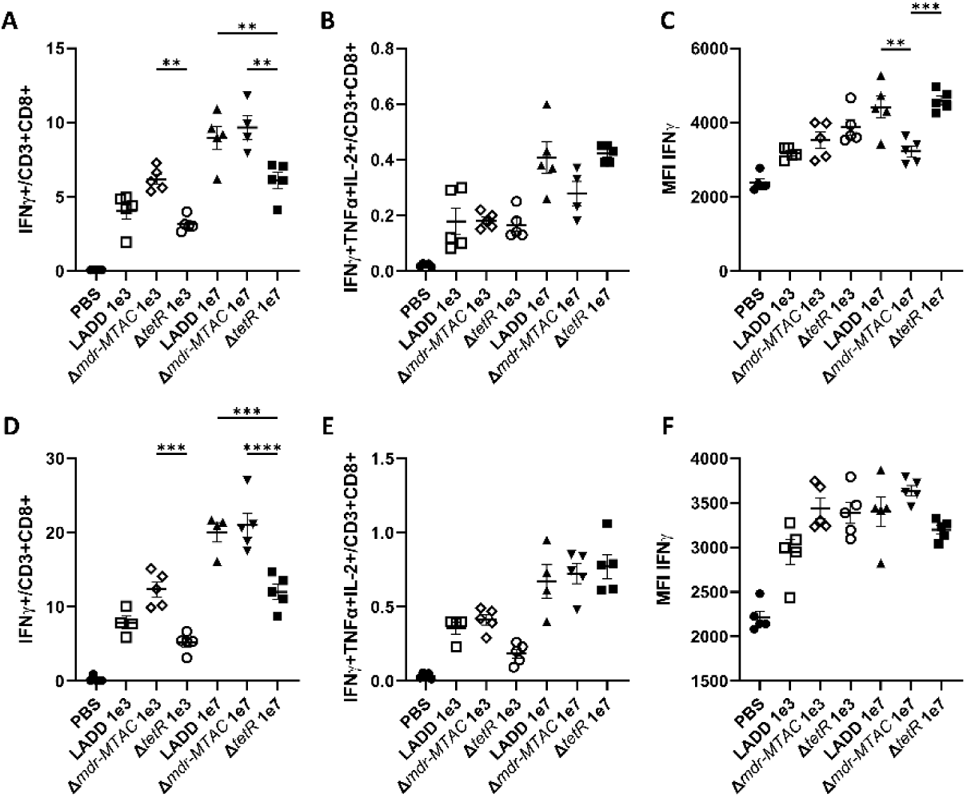
OVA and B8R-specific antigen responses in T-cell response and IFNγ expression for peptide stimulated splenocytes from mice immunized with type I Interferon altered vaccine strains (goes with figure 5). Mice were immunized with 1×10^3^ the indicated strain of *L. monocytogenes* expressing B8R and OVA. OVA specific T-cell responses were analyzed by *ex vivo* peptide stimulation at day 7 post immunization for CD8+ T-cells producing A) IFNγ, and B) IFNγ+TNFα and IL-2+, as well as C) IFNγ expression. B8R specific T-cell responses were analyzed by *ex vivo* peptide stimulation at day 7 post immunization for CD8+ T-cells producing D) IFNγ, and E) IFNγ+TNFα and IL-2+, as well as F) IFNγ expression. Data are representative of two independent experiments of 4-5 mice per group Significance was determined by a one-way ANOVA with Tukey’s correction. * p < 0.05.

**Supplemental figure 9 (goes with figure 5) -.**
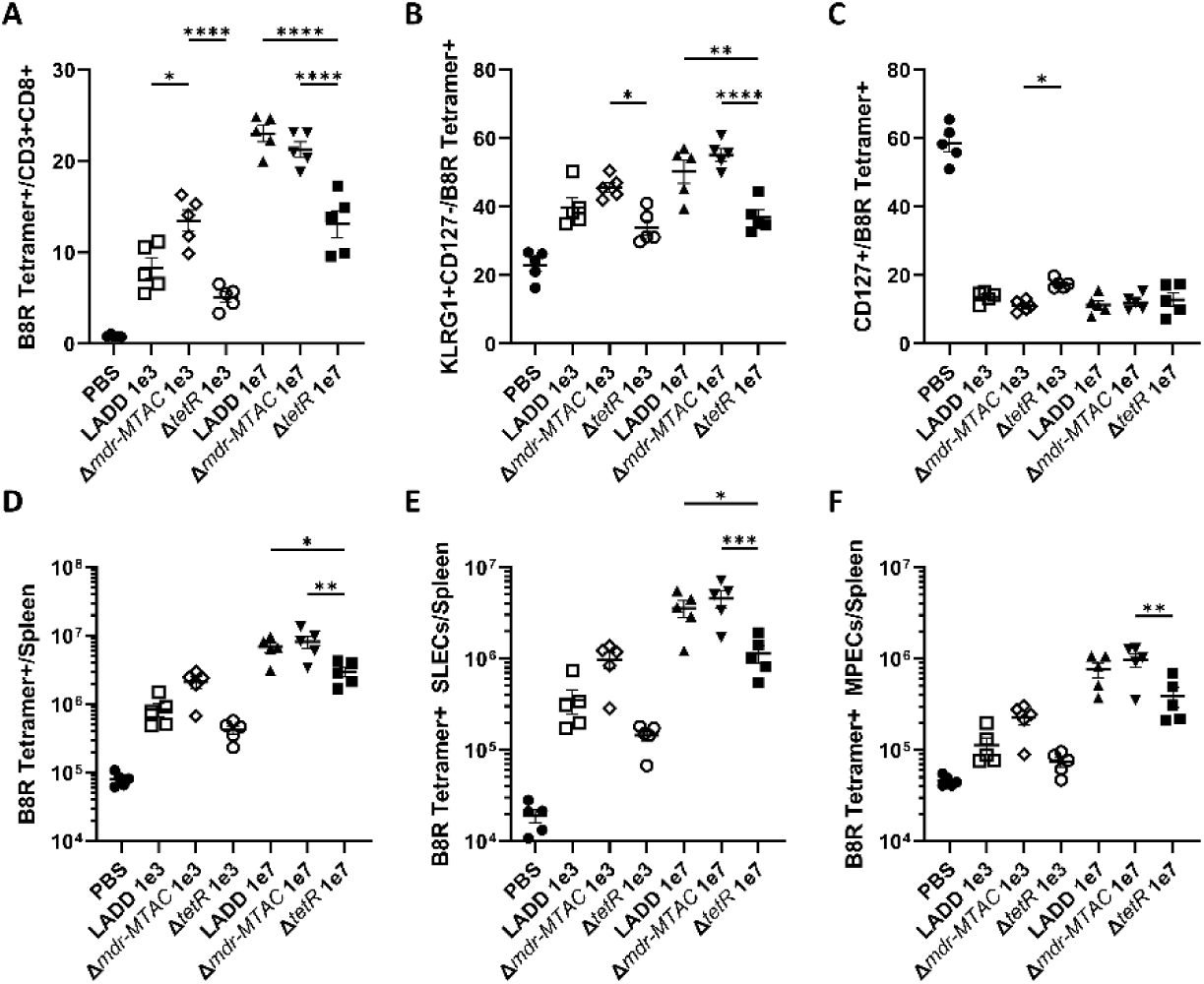
The B8R-specific T-cell response is enhanced in mice immunized with type I Interferon hypo-inducing *L. monocytogenes.* Mice were immunized with indicated strain and dose of *L. monocytogenes* and B8R-specific T-cell A) frequencies and D) numbers per spleen, or B8R-specific SLEC B) frequencies and D) numbers per spleen, or B8R-specific MPEC C) frequencies and F) numbers per spleen, were assessed. Data shown are representative of two independent experiments of 4-5 mice per group. Significance was determined by a one-way ANOVA with Tukey’s correction. * p < 0.05.

**Supplemental figure 10 (goes with figure 6) –.**
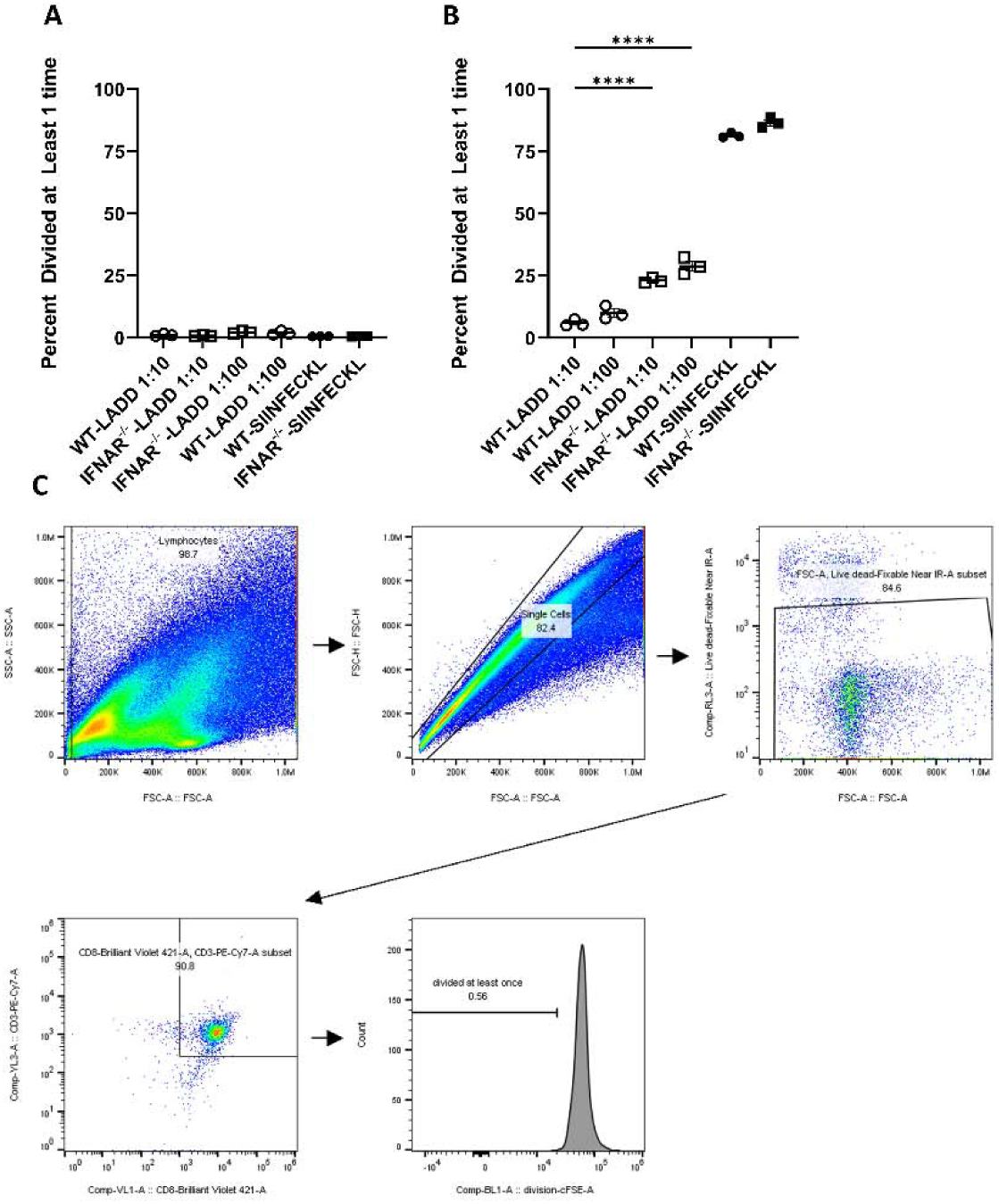
IFNAR^−/−^ APCs promote enhanced T-cell proliferation in *ex vivo* T-cell proliferation assays. CD11c+ splenocytes from B16-Flt3L tumor-bearing WT or IFNAR^−/−^ mice were isolated and infected with *L. monocytogenes* expressing OVA and B8R for 6 hours prior to the addition of CFSE-labeled OT-I T-cells. The percentage of proliferated CD3+CD8+ T-cells was determined by flow cytometry for CFSE dilution at A) 24 or B) 48 hours post addition of CFSE-lavedl OT-I T-cells. Data shown are representative of two independent experiments. Significance was determined by a one-way ANOVA with Bonferroni’s correction. * p < 0.05.

